# Confocal Laser-Scanning Fluorescence-Lifetime Single-Molecule Localisation Microscopy

**DOI:** 10.1101/2020.08.25.266387

**Authors:** Jan Christoph Thiele, Dominic Helmerich, Nazar Oleksiievets, Roman Tsukanov, Eugenia Butkevich, Markus Sauer, Oleksii Nevskyi, Jörg Enderlein

**Affiliations:** III. Institute of Physics – Biophysics, Georg August University, Göttingen, Germany; Department of Biotechnology and Biophysics, Biocenter, University of Würzburg, Am Hubland, 97074 Würzburg, Germany; Cluster of Excellence “Multiscale Bioimaging: from Molecular Machines to Networks of Excitable Cells” (MBExC), Georg August University, Göttingen, Germany

## Abstract

Fluorescence lifetime imaging microscopy (FLIM) is an important technique that adds another dimension to the intensity and colour information of conventional microscopy. In particular, it allows for multiplexing fluorescent labels that have otherwise similar spectral properties. Currently, the only super-resolution technique that is capable of recording super-resolved images with lifetime information is STimulated Emission Depletion (STED) microscopy. In contrast, all Single-Molecule Localisation Microscopy (SMLM) techniques that employ wide-field cameras completely lack the lifetime dimension. Here, we combine Fluorescence-Lifetime Confocal Laser-Scanning Microscopy (FL-CLSM) with SMLM for realising single-molecule localisation-based fluorescence-lifetime super-resolution imaging (FL-SMLM). Besides yielding images with a spatial resolution much beyond the diffraction limit, it determines the fluorescence lifetime of all localised molecules. We validate our technique by applying it to direct STochastic Optical Reconstruction Microscopy (dSTORM) and Points Accumulation for Imaging in Nanoscale Topography (PAINT) imaging of fixed cells, and we demonstrate its multiplexing capability on samples with two different labels that differ only by fluorescence lifetime but not by their spectral properties.

## Introduction

Confocal Laser-Scanning Microscopy (CLSM) is one of the most important microscopy techniques for biology and medicine. Its fundamental purpose is to provide so-called optical sectioning and thus to enable the recording of three-dimensional images, which is impossible to achieve with conventional wide-field microscopy. Its disadvantage, when compared to wide-field microscopy, is its inherently slow image acquisition speed, because image formation is realised by sequentially scanning single or multiple foci over a sample. This limits also its overall light throughput (small dwell time per scan position), which is one reason why CLSM was nearly never used for single-molecule localisation based super-resolution microscopy (Single Molecule Localisation Microscopy or SMLM), such as Photo-activation Localisation Microscopy (PALM),(*1*) (direct) Stochastic Optical Reconstruction Microscopy (dSTORM),(*2, 3*) or Points Accumulation for Imaging in Nanoscale Topography (PAINT).(*4, 5*) There are only two exceptions,(*6, 7*) one where a spinning-disk CLSM was employed for PAINT, exploiting the superior out-of-plane light rejection of a CLSM that is so important for reducing background from freely diffusing dyes in PAINT, and one where a spinning-disk CLSM was employed for STORM with self-blinking dyes, where it was used for reducing excitation intensity. Besides efficient out-of-plane signal rejection, which also enhances contrast and facilitates deep-tissue imaging,(*8*) CLSM offers several additional advantages that should make it attractive for SMLM. Firstly, single-focus CLSM uses single-point detectors which can be operated in single-photon counting mode (Geiger mode) and thus provide shot-noise limited detection, in contrast to emCCD or sCMOS cameras used in conventional wide-field SMLM, with their read-out, thermal, and electronic noise. Secondly, when using Geiger mode for light detection, CLSM records the positions of single-photon detection events in a quasi-continuous, non-pixelated way, thus avoiding pixel size to affect single-molecule localisation accuracy.(*9*) Thirdly, and most interestingly, it allows for measuring fluorescence lifetimes, thus allowing to combine Fluorescence Life-time Imaging Microscopy (FLIM) with SMLM. This offers, for example, the option to co-localise different molecular species that differ only by their lifetime while having similar excitation and emission spectra,(*10*) thus efficiently circumventing all problems connected with chromatic aberration that trouble so much multi-colour SMLM.(*11, 12*) Especially for state-of-the-art SMLM, which now routinely achieves a lateral resolution of only a few nanometers, chromatic aberration is a serious issue,(*13*) in particular when trying to study biological interactions or the relative arrangement of different cellular structures with respect to each other.

Several solutions to the chromatic aberration problem have been proposed in the past. Recently, an aberration-free multi-colour method of SMLM called spectral-demixing dSTORM was presented.(*14*) This method works well for fluorophores showing good switching performance while utilising the same imaging buffer. The fluorescence signal of the different molecules is separated spectrally, and ratiometric fluorescence measurements are used for spectral demixing and (co-)localising different kinds of molecules. One step further in this direction was the implementation of spectrally-resolved SMLM, where full spectra are measured and used for sorting different molecules and their localisations.(*15*) A very fascinating approach is multi-colour SMLM that combines PSF engineering with deep learning for identifying and sorting different molecular species without the need of spectrally resolved imaging.(*16*) In frequency-based multiplexing STORM/DNA-PAINT,(*17*) one uses frequency-encoded multiplexed excitation and colour-blind detection to circumvent chromatic-aberration problems. Another clever solution is Exchange-PAINT,(*18*) which sequentially images different targets with the same dye but uses different DNA-tags for directing the dye to different targets decorated with complementary DNA-strands. Similarly, barcoding PAINT(*19*) exploits the different binding kinetics of imager and docking strands for distinguishing between different target sites. Because one uses the same dye for all the different structures, chromatic aberrations do not impact the SMLM results, but the prize is increasing image acquisition time, ca. linearly increasing with the number of different targets one wants to resolve. Finally, the recently introduced MINFLUX(*20*) allows for super-resolution imaging with unprecedented accuracy of only a few nanometers and can be used for chromatic-aberration free multi-colour imaging.(*21*) Similar to the confocal laser-scanning SMLM that will be presented here, it is also based on scanning, but in an asynchronous manner, so that it can currently localise only one individual molecule at any time.

In this work, we present the first (to our knowledge) realisation of SMLM with a time-resolved CLSM using single-photon avalanche-diodes (SPADs) for detection, and a rapid laser-scanning unit for excitation beam scanning. This unit enables us to record images with reasonable acquisition speed as required for efficient SMLM. Our approach combines all the advantages of CLSM with those of SMLM: axial sectioning, shot-noise limited single-photon detection, pixel-free continuous position data, and fluorescence lifetime information of CLSM with the exceptional spatial resolution and single-molecule identification of SMLM. We first demonstrate the feasibility of using CLSM for fluorescence lifetime SMLM (FL-SMLM) by imaging labelled fixed cell samples by combining CLSM with two of the most widely used variants of SMLM, dSTORM (for imaging microtubules and clathrin in human mesenchymal stem cells) and DNA-PAINT (for imaging cellular chromatin in COS-7 cells). To demonstrate the fluorescence lifetime multiplexing capability of FL-SMLM, we record images of polymer beads that are surface-labelled with two different dyes and two cellular targets (microtubules and clathrin in COS7 cells). Our results show that confocal laser-scanning FL-SMLM has great potential for many applications, extending the dimensions of fluorescence super-resolution microscopy by fluorescence lifetime.

## Results and Discussion

Confocal laser-scanning SMLM measurements were carried out on a custom-built time-resolved confocal microscope equipped with a fast laser scanner, see Fig.1a (for more details see SI). For dSTORM measurements, a region of interest of 10 µm × 10 µm was scanned with an image scan rate of ∼27 Hz. Excitation was done with a pulsed laser at 640 nm wavelength, a repetition rate of 40 MHz, and a pulse width of ∼50 ps. Single fluorescence photons were detected with a single-photon avalanche diode (SPAD), and photon detection events were correlated in time to excitation pulses (Time-Correlated Single-Photon Counting or TCSPC) with high-speed electronics. For data analysis, recorded photons were converted into a stack of intensity images, always combining 10 subsequent scans into one image to minimise distortions by those labels that are switching while being scanned. The stack of intensity images is then used to localise single molecules and to identify switching events. These localisations were subsequently used to reconstruct a super-resolved image (similar to conventional wide-field dSTORM). Taking full advantage of our TCSPC detection, fluorescence lifetimes of each localised molecule were determined by pooling all photons associated with it and fitting the resulting lifetime histogram with a mono-exponential decay function. To increase the number of photons per localisation, identical localisations in subsequent frames were merged. Using this lifetime information, super-resolved fluorescence lifetime images were reconstructed.

**Fig. 1.**
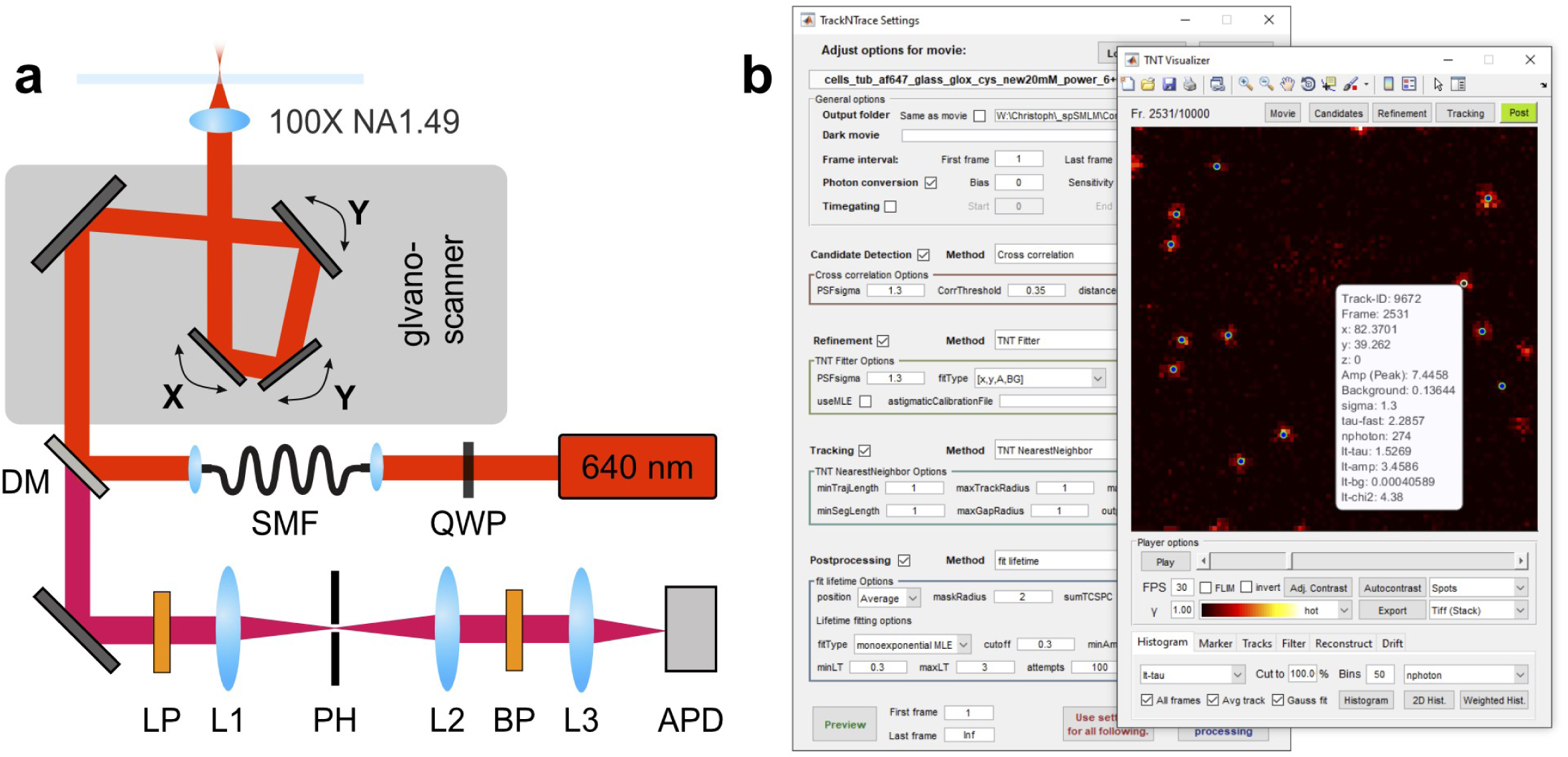
Measurement and data evaluation. **(a)** Schema of the confocal setup: The pulsed640 nm excitation light is converted to radial polarisation with a quarter wave plate (QWP), passes through a single mode fibre (SMF), is reflected by a dichroic mirror (DM) into a galvometeric laser scanner and focused by the objective. The collected fluorescent emission from the sample is descanned, passes the DM and is focused on the pinhole (PH), then on single-photon detector (APD) using the lenses (L1, L2 and L3). The long pass filter (LP) and the band pass filter (BP) are blocking scattered excitation light. **(b)** The scanning SMLM data analysis from raw data to reconstruction was done in an extended version of TrackNTrace.

To make the data analysis for confocal laser-scanning FL-SMLM widely available, the complete data processing pipeline was integrated into a Matlab-based GUI app, see Fig.1b. This app builds on the existing open-source framework TrackNTrace(*22*) and was now extended for processing also TCSPC data. The app supports FLIM data and comes with a dedicated plugin that pools photon detections for each localised molecule and executes lifetime fits. The data visualiser offers filtering of localisations, drift correction with Redundant Cross-Correlation (RCC),(*23*) and reconstruction of super-resolved lifetime images. More details can be found in the SI.

To demonstrate the applicability of confocal laser-scanning SMLM for biological samples, we performed dSTORM of microtubules in fixed immunolabelled hMSC cells. Alexa 647 is a benchmark dye for dSTORM due to its outstanding photostability, brightness, and optimised blinking behaviour.(*24*) The key parameters here are the high number of photons per switching event and the possibility to tune the switching kinetics through the composition of the imaging buffer and a suitable adjustment of excitation power. Conventional CLSM and confocal laser-scanning dSTORM images are presented in Fig. 2b. A Gaussian fit of the fluorescence lifetime histogram obtained from the fitted lifetime values of all identified molecules in the region of interest gives a mean value of 1.51 ns for the fluorescence lifetime of Alexa 647-labeled antibodies, which is in agreement with literature data.(*25*) Moreover, the width of the obtained lifetime distribution (see Fig. 2c) is close to being shotnoise limited, so that multiplexing by fluorescence lifetime seems a very promising prospect for FL-SMLM. The average number of detected photons per switching cycle was calculated to be 1085 photons/event (see Fig. S4a), which is lower than for imaging with a wide-field setup.(*24*) We attribute this lower photon count to light losses in the detection pathway (due to dichroic mirrors, pinhole and emission filters). To better compare the performance of confocal laser-scanning dSTORM with conventional wide-field dSTORM, we imaged the same sample that we used for FL-SMLM with a custom-built wide-field setup. The comparison is shown in Fig. S2. To quantify the relative performance of both techniques, we determined single microtubule cross-sections, and we estimated their diameter to be 64 nm FWHM for confocal laser-scanning dSTORM, and 53 nm FWHM for conventional wide-field dSTORM. The apparent size of microtubules in both cases is larger than their actual value, which we attribute to the extra size of the secondary immunolabels. We also calculated Fourier Ring Correlation (FRC) maps using the NanoJ-SQUIRREL plugin (see Fig. S1).(*26*) and found that the average resolution for both images was 50 nm and 48 nm for confocal laser-scanning dSTORM and conventional wide-field dSTORM, respectively. The localisation precision was estimated following modified Mortensen’s equation,(*9, 27, 28*) and an average value was found to be 8.6 nm and 9.2 nm, respectively. In summary, we find that both approaches show a similar performance, which demonstrates that confocal laser-scanning dSTORM can be a promising and versatile super-resolution technique adding the important fluorescence lifetime dimension to the picture. Furthermore, the sectioning capabilities of the confocal imaging system offers the possibility to image dense 3D structures. To illustrate this, we have recorded *z*-stacks of images of Alexa 647 labelled tubulin in COS7 cells (see Fig. S2)

**Fig. 2.**
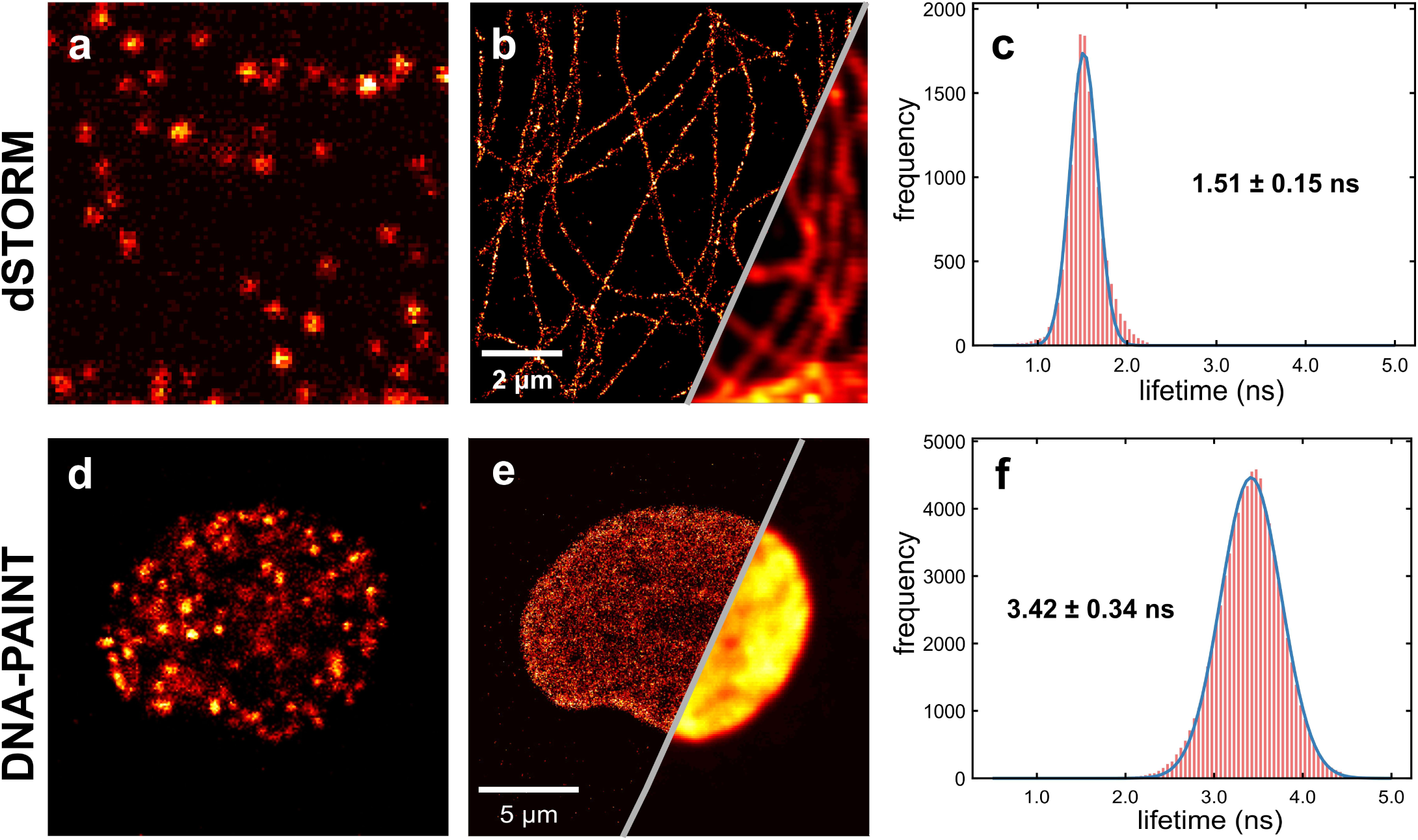
Confocal dSTORM and DNA-PAINT imaging in cells. **(a-c)** Confocal laser-scanning dSTORM images of tubulin filaments in hMS cells labelled with Alexa 647. **(d-f)** Confocal laser-scanning DNA-PAINT images of cellular chromatin in COS-7 cells utilising DNA-labelled ATTO 655. **(a**,**d)** Example of a single frame during acquisition. **(b**,**e)** Corresponding diffraction-limited and super-resolved images. **(c**,**f)** Intensity-weighted lifetime histograms, based on individual single-molecule localisations.

We applied our approach also to perform confocal laser-scanning DNA-PAINT. DNA-PAINT is a recently developed and highly promising alternative to dSTORM, because it circumvents the inherent photobleaching limitations of dSTORM by employing dye-labelled imager DNA-strands that reversibly bind to targets of interest that are themselves labelled with complementary target DNA-strands. For DNA-PAINT (and PAINT in general), optical sectioning is critical for efficiently suppressing fluorescent background from freely diffusing imager strands. To demonstrate confocal laser-scanning DNA-PAINT, we imaged histone H2B in COS-7 cells (Fig. 2 d-f) that was fused to mTagBFP which was subsequently high-affinity labelled with DNA docking strands using FluoTag®-Q anti-TagBFP nanobodies.(*29*) Using nanobodies for labelling minimises the distance between dye and target, thus significantly reducing so called linkage errors and respectively increasing localisation precision. ATTO 655 was used for DNA-PAINT because of its high brightness and low unspecific binding to both coverslip surface and cell organelles. Although we reduced the concentration of imager strands, as compared to conventional DNA-PAINT,(*18*) by an order of magnitude (to 0.25 nM), we could register a sufficiently large number of single-molecule localisations due to the dense packing of histone targets inside the nucleus. A typical single frame from a recorded movie is shown in Fig. 2d, and Fig. 2e presents a comparison between super-resolved PAINT and conventional CLSM. The average localisation precision of the reconstructed confocal laser-scanning DNA-PAINT image was found to be 18 nm. For more detailed information and comparison with conventional DNA-PAINT see Fig. S3. The corresponding lifetime distribution for all localised molecules is shown in Fig. 2f. There, we find a lifetime value for ATTO 655 that is longer than the reported value for free dye, which we attribute to its conjugation to single-stranded imager DNA.

To demonstrate the sectioning as well as multiplexing capabilities of confocal laser-scanning SMLM, we imaged polymer beads (d = 3 µm) labelled with two different fluorophores (Alexa 647 and ATTO 655) bound to DNA. On our wide-field microscope, we could not detect single switching events when focusing at the centre of the beads. In contrast, it was possible to localise switching molecules (Fig. 3a) with the CLSM, to determine their fluorescence lifetimes, and to reconstruct a fluorescence-lifetime dSTORM image (Fig. 3b). The image shows two beads that are labelled with two different dyes, namely Alexa 647 having an average fluorescent lifetime of 1.4 ns and ATTO 655 with 2.4 ns. Details on data processing and the resulting lifetime histogram can be found in the methods section (see Fig. S5).

**Fig. 3.**
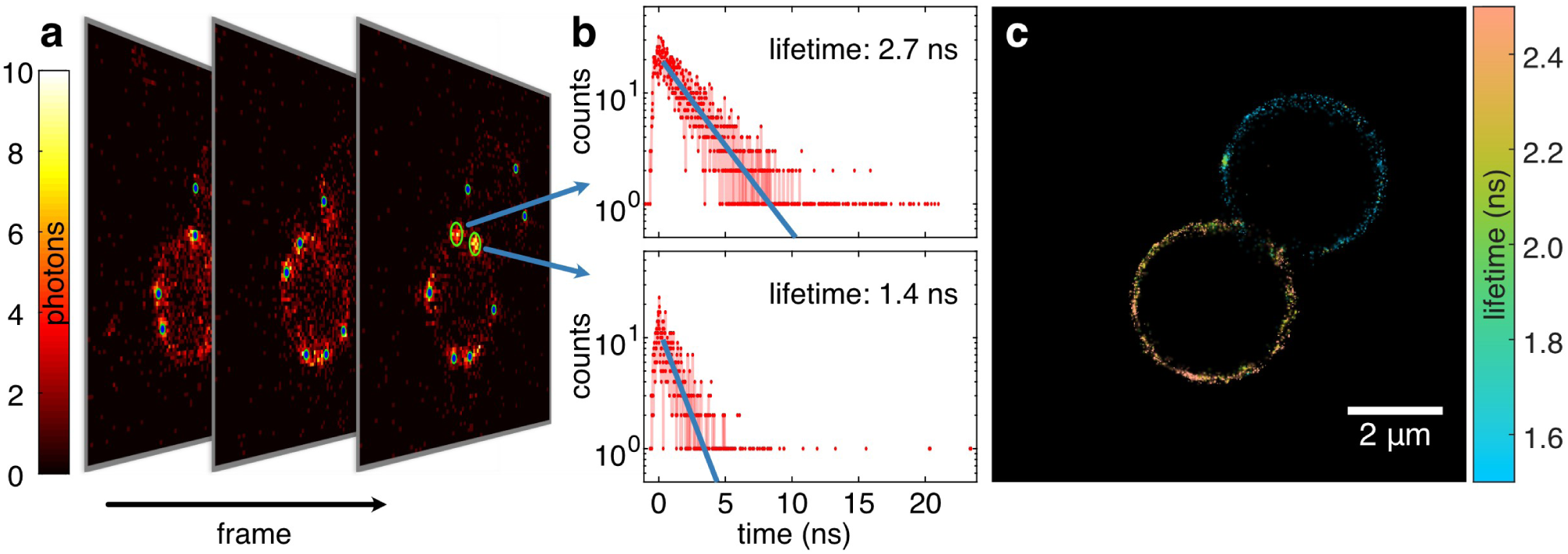
Lifetime-based multiplexed dSTORM imaging of polymer beads labelled with two different dyes. **(a)** Typical frame from a recorded movie including single-molecule localisations. **(b)** Histogram of photon arrival times (TCSPC) for two indicated localisations. Lifetime is determined with a mono-exponential fit (blue line). **(c)** Super-resolved image reconstruction including lifetime information. The two different lifetime values for molecules on both beads reveal that the beads are labelled with different fluorophores (Alexa 647 and ATTO 655).

To enable fluorescence-lifetime multiplexing dSTORM, it is crucial to use buffer conditions which are compatible with all fluorophores. This includes control of sufficient on- and off-switching rates while guaranteeing high dye brightness. Typically, a primary thiol is used to enhance off-switching rates. The standard buffer for Alexa 647 additionally contains GLOX to create an oxygen-depleted environment which reduces permanent photobleaching. We found that an oxygen-depleted environment results in low on-switching rates for ATTO 655, probably due to a long-living non-fluorescent reduced form.(*30*) However, using a buffer that only contains thiol was sufficient to dSTORM both fluorophores.

To validate fluorescence-lifetime multiplexing SMLM on biological samples, we performed dSTORM imaging of Alexa 647 labelled β-tubulin and ATTO 655 labelled clathrin in COS7 cells. Thus, two different targets are labelled with two spectroscopically similar dyes with different fluorescence lifetimes. To improve lifetime contrast and brightness of both dyes all dual-label dSTORM cell images were done in D_2_O instead of PBS buffer and with the same thiol concentration as for the bead imaging.(*31, 32*) Additionally, a short PEG-4 linker was inserted between the secondary antibody and ATTO 655 to reduce interactions influencing its lifetime (see methods section for more details). To classify localisations, pattern matching was used as an alternative to lifetime fitting. For this, the probability that a reference TCSPC decay curve (see Fig. 4a) produces that of a localised molecule is calculated and the molecule classified according to the reference that yields the highest probability.

**Fig. 4.**
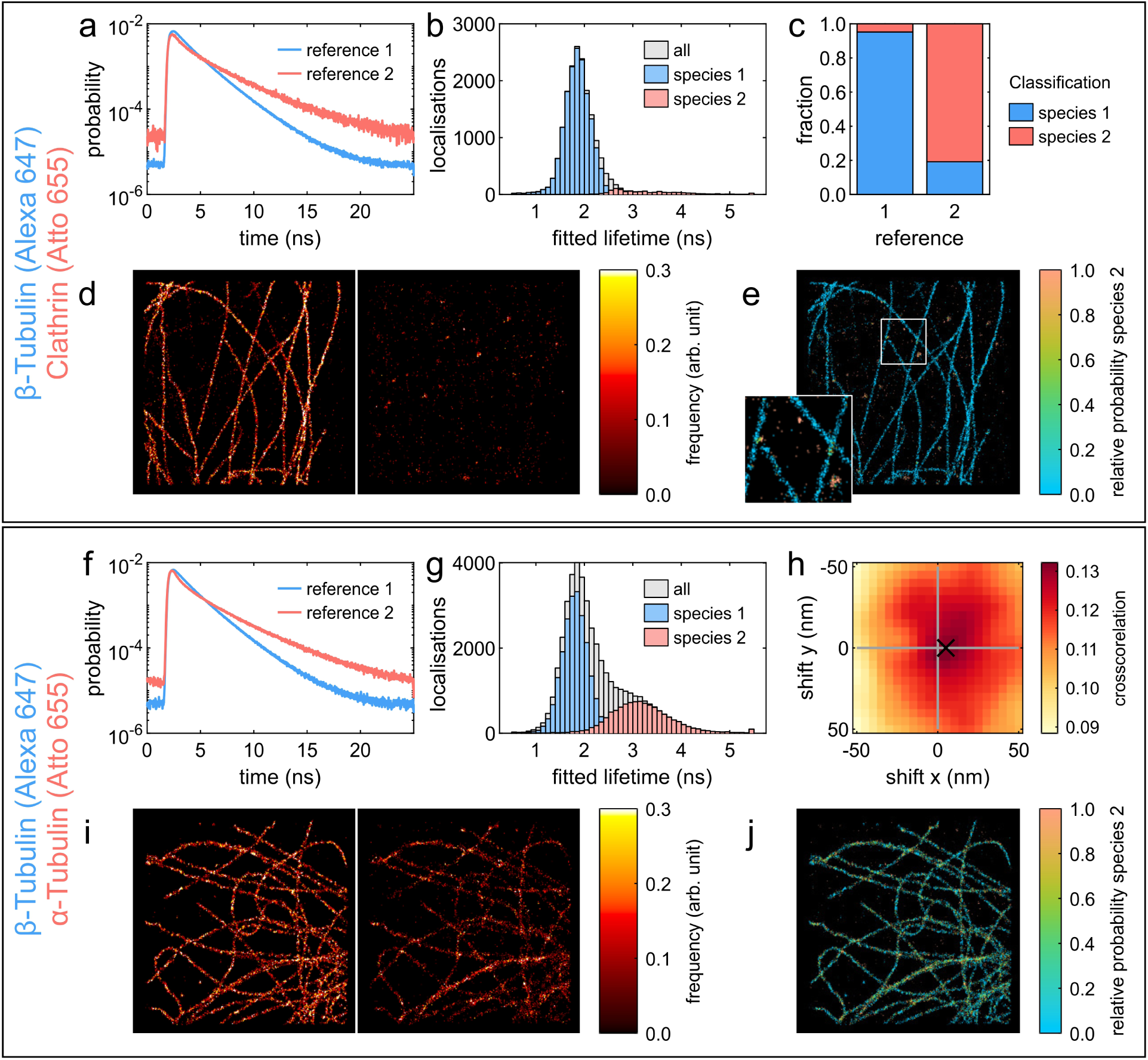
Lifetime-based multiplexed dSTORM imaging in fixed COS7 cells. All dSTORM images are 10×10 µm and reconstructed with a 10 nm Gaussian PSF. **(a**,**f)** Reference TCSPCs measured from samples labelled only with one dye (reference 1 – Alexa 647, reference 2 – ATO 655). **(b, g)** Fitted lifetime histograms based on individual single-molecule localisations for Alexa 647 β-tubulin / ATTO 655 clathrin labelled and Alexa 647 β-tubulin / ATTO 655 α-tubulin labelled samples, correspondingly. *Species 1* represents Alexa 647, *species 2* represents ATTO 655 dye. **(c)** Calculated cross-talks for the wrong species assignment by classify the reference samples. *Species 1* represents Alexa 647, *species 2* represents ATTO 655 dye. **(d, i)** Super-resolved confocal dSTORM images of localisations classified (d) Alexa 647 β-tubulin (left) / ATTO 655 clathrin (right) and (i) Alexa 647 β-tubulin (left) / ATTO 655 α-tubulin (right). **(e**,**j)** Super-resolved probability images obtained by pattern matching analysis. *Species 2* represents ATTO 655. The intensity was exponentiated with a γ of 0.7. **(h)** Cross-correlation calculated between super-resolved images of *species 1* and *species 2* to estimate possible chromatic aberration shift. The underlying images were reconstructed with 5 nm pixel and a 5 nm Gaussian PSF.

This has several advantages: it uses all the detected photons, it is not assuming that the fluorescence decay is mono-exponential, and it is fit-free and therefore fast and stable. From Fig. 4b it can be seen that the lifetime distributions of both species overlap. Nevertheless, it is possible to identify species 1 with less than 5 % false classifications and species 2 with less than 20 %, as shown Fig. 4c, by rejecting classifications that have a relative probability below 99 % (see methods). Corresponding dSTORM images for both targets are shown in Fig. 4d. The resulting super-resolved probability image based on the pattern matching analysis is shown in Fig. 4e.

One of the biggest advantages of confocal scanning SMLM is the possibility to avoid chromatic aberration artefacts in colocalisation measurements. To demonstrate this, we labelled α-tubulin and β-tubulin, which are colocalised in microtubules, with ATTO 655 and Alexa 647 correspondingly. For the probability analysis, corresponding reference TCSPC curves were measured (see Fig. 4f) of only ATTO 655 labelled α-tubulin and only Alexa 647 labelled β-tubulin samples. To verify correct colocalization between the two species, we calculated the cross-correlation (shown in Fig. 4h) between corresponding images (Fig. 4i). Its maximum at shift value of 5 nm indicates that there is a negligible shift between the channels. This can be also seen in the super-resolved probability image presented in Fig. 4j.

In summary, we have presented confocal laser-scanning FL-SMLM which combines sectioning and lifetime information with super-resolution imaging. The technique is straight-forward to implement on a commercial CLSM with TCSPC capability and fast laser scanning. As light exposure is limited only to the scanned area, it is possible to sequentially image different regions of interest without prior photobleaching of other regions. We demonstrated FL-SMLM both with dSTORM and DNA-PAINT, which are the most commonly used SMLM modalities. The high lifetime resolution enables lifetime-based multiplexing within the same spectral window, distinguishing different fluorescent labels solely by their lifetimes. In combination with the optical sectioning of a CLSM, this allows for chromatic aberration-free super-resolution imaging of multiple cellular structures. The additional lifetime information in FL-SMLM offers also the fascinating prospect of lifetime-based super-resolution FRET imaging.(*33*) Another potential application is to use the lifetime information for combining lateral super-resolution of SMLM with the superior axial super-resolution of Metal-Induced Energy Transfer (MIET) imaging.(*34, 35*) This could enable 3D super-resolution imaging with exceptionally high isotropic resolution for potential applications in structural biology.

## Materials and Methods

### Confocal microscopy

Fluorescence lifetime measurements were performed on a custom-built confocal setup. For the xcitation a 640 nm 40 MHz pulsed diode laser (PDL 800-B driver with LDH-D-C-640 diode, PicoQuant) was utilised. A quarter-wave-plate was used in the excitation path to convert linear to circular polarisation. The laser beam was coupled into a single-mode fiber (PMC-460Si-3.0-NA012-3APC-150-P, Schäfter + Kirchhoff) with a fiber-coupler (60SMS-1-4-RGBV-11-47, Schäfter + Kirchhoff). After the fiber, the output beam was collimated by an air objective (UPlanSApo 10× /0.40 NA, Olympus). After passing thought a clean-up filter (MaxDiode 640/8, Semrock), an ultra-flat quad-band dichroic mirror (ZT405/488/561/640rpc, Chroma) was used to direct the excitation light into the microscope. The excitation beam was directed into a laser scanning system (FLIMbee, PicoQuant) and then into a custom side port of the microscope (IX73, Olympus). The three galvo mirrors in the scanning system are deflecting the beam while preserving the beam position in the backfocal plane of the objective (UApo N 100× /1.49 NA oil, Olympus). The sample position can be adjusted a using manual XY stage (Olympus) and a z-piezo stage (Nano-ZL100, MadCityLabs). Emission fluorescence light was collected by the same objective and de-scanned in the scanning system. Afterwards, the achromatic lens (TTL180-A, Thorlabs) was used to focus the beam onto the pinhole (100 µm P100S, Thorlabs). The excitation laser light was blocked in the emission path by a long-pass filter (647 LP Edge Basic, Semrock). Then, the emission light was collimated by a 100 mm lens. A band-pass filter (BrightLine HC 692/40, Semrock) was used to reject scattered excitation light. Finally, the emission light was focused onto a SPAD-detector (SPCM-AQRH, Excelitas) with an achromatic lens (AC254-030-A-ML, Thorlabs). The output signal of the photon detector was recorded by a TCSPC system (HydraHarp 400, PicoQuant) which was synchronised with the triggering signal from the excitation laser. Measurements were acquired with the software (SymPho-Time 64, PicoQuant), which controlled both the TCSPC system and the scanner system. Typically, sample scan with a virtual pixel size of 100 nm, and a dwell time of 2.5 µs/pixel and a TCSPC time resolution of 16 ps were chosen.

To evaluate the performance of the laser scanner, TetraSpeck microspheres were measured with the same parameters as used in the dSTORM measurements (10×10 µm region of interest, 100 nm pixel size and 2.5 µs dwell time). For the evaluation, 10 scans were binned to one frame as it was done for the dSTORM measurement. For the analysis TrackNTrace was used to independently localise the spheres in each frame. The lateral drift was corrected by subtracting the moving average over 100 frames of the centre of gravity of all localisations. Fig. S7 shows the sample (a) and an exemplary track (b). The average standard deviation of the tracks is 4.0 nm in *x* and 5.2 nm in *y*. It is remarkable that the standard deviation along the fast scanning axis (*x*) is slightly lower than along the slow axis (*y*).

### Wide-field microscopy

Measurements were performed using a custom-built optical setup, as described previously.(*29*) The excitation was performed using a 638 nm continuous-wave laser (PhoxX+ 638-150, Omicron). The laser beam was coupled into a single-mode optical fiber (P1-460B-FC-2, Thorlabs) with a typical coupling efficiency of 50 percent. The collimated laser beam was expanded by a factor of ∼3.6× using a telescope lenses after the fiber. The beam was focused onto the back focal plane of the TIRF objective (UAPON 100x oil, 1.49 NA, Olympus) using achromatic lens (f=200 mm, AC508-200-A-ML, Thorlabs). Switching between the EPI and Total Internal Reflection (TIR) illumination schemes was achieved by a mechanical shifting the beam with respect to the objective lens using the translation stage (LNR50M, Thorlabs). Fluorescence emission light was collected using the same objective lens. Samples were placed onto the XY translation stage (M-406, Newport). An independent one-dimensional translation stage (LNR25/M, Thorlabs) was equipped with differential micrometer screw (DRV3, Thorlabs) for focusing by moving the objective lens. Emission fluorescence light was spectrally decoupled from the scattered excitation laser light using a multi-band dichroic mirror (Di03 R405/488/532/635, Semrock) and a band-pass filter (BrightLine HC 692/40, Semrock). Using tube lens (AC254-200-A-ML, Thorlabs), imaging plane was formed on an adjustable slit aperture (SP60, OWIS). The latter was employed to select a region of interest within the field of view. Two Lenses (AC254-100-A, Thorlabs) and (AC508-150-A-ML, Thorlabs) were used to form the image the on an emCCD camera (iXon Ultra 897, Andor). The total magnification for imaging was 166.6X, resulting in an effective pixel size in sample space of 103.5 nm. Image stacks of 20000 frames were acquired with an exposure time of 10 ms. The recorded images then were analysed with the ImageJ plugin (ThunderSTORM)(*36*) for determining the positions of single emitters. In all the experiments, the same localisation parameters were used for super-resolution image reconstruction. Localisations containing more than 500 photons and less than 5000 photons were taken into account.

### dSTORM imaging

For conventional/confocal dSTORM imaging utilizing only Alexa 647 dye, an enzymatic oxygen scavenging buffer (glucose oxidase 0.5 mg/mL, catalase 40 g/mL, glucose 10% w/v in PBS pH 7.4) with addition of thiol (20 mM Cysteamine) was used. For multiplexed confocal dSTORM imaging of beads / fixed cells the switching buffer containing only thiol (5 mM Cysteamine) in PBS and in D_2_O was used, correspondingly.

### Points accumulation for imaging in nanoscale topography (PAINT) imaging

The DNA docking strands (Biomers GmbH, Ulm, Germany) were functionalized with an azide group at 5’-end. The coupling of the docking strands to the unconjugated nanobodies FluoTag®-Q anti-TagBFP (NanoTag Biotechnologies GmbH, Göttingen, Germany, Cat. No: N0501) was performed according to the procedure described by Schlichthärle and colleagues.(*37*) The imager strand (Eurofins Genomics) was labelled with ATTO 655 fluorophore at the 3’-end. It was aliquoted in TE buffer (Tris 10 mM, EDTA 1 mM, pH 8.0) at a concentration of 1 µM and stored at -20° C. Prior to the experiment, the strands were diluted to the final concentration of 0.25 nM in PBS buffer, containing 500 mM NaCl. Imager strand solution in PBS buffer (500 µL) were added into the well and incubated for 5 min before the acquisition. The following acquisition settings were used: area 20×20 µm, virtual pixel size 100 nm, dwell time 2.5 µs/pixel and a TCSPC time resolution 16 ps. 10 scans were combined together as for the dSTORM measurements and analysed by TrackNTraceLT software. The Fourier ring correlation (FRC) maps were obtained for reconstructed super-resolved images (see Figure S3) and the average resolutions were calculated 45 nm and 41 nm for confocal DNA-PAINT and conventional DNA-PAINT, respectively. The following average localization precision values were calculated 18 nm and 26 nm, correspondingly. The measured lifetimes values were as following: non-conjugated ATTO 655 dye in solution 1.83±0.01 ns, DNA-dye construct in solution 2.69 ± 0.01 ns and DNA-dye construct attached to its complementary docking strand inside the nucleus 3.44 ± 0.3 ns. As expected, the lifetimes of freely diffusing fluorophore is shorter than DNA-conjugated dye due to the DNA-dye interactions.

### TrackNTrace Lifetime Edition

In order to facilitate the data analysis of scanning confocal super-resolution imaging, we developed an extended version of the Matlab based framework TrackNTrace.(*22*) To allow the processing of FLIM data the image import was generalised to use file type specific plugins. The existing support for TIFF files was moved to a dedicated import plugin and a plugin in for PicoQuant Time Tagged File Format (PTU) was implemented. Since the PTU format stores a stream of photons with relative time information and position markers it is not possible to read specific photons. To allow random access to photon data based on the frame and position they were detected at, an index of all photons including their position, frame, micro time (TCSPC bin) and channel was created and stored as MAT-file, which is hdf5-based and allows random access. During the creation of the photon index scanning artefacts where corrected as described below. The photon index was used to generate a stack of intensity images, allowing a user-defined frame binning and time-gating. The intensity images were used to perform the single molecule detection, refinement and tracking with the plugins provided by the original version of TrackAndTrace. Subsequently, a fourth step was added, the post-processing of the localisations. For this step a plugin to determine the fluorescence lifetime of the localisations was created. First, the localisations can be refined by either averaging the position the position over all frames of the track or by refitting the localisation in a sum image of all frames of the track. Then, the photons within a predefined radius (default 2σ_PSF_) around each localisation are extracted from the photon index to generate the TCSPC histograms of the localisations. To determine the lifetime each TCSPC histograms is fitted with a mono-exponential tailfit using a Nelder–Mead simplex algorithm to minimise the negative log-likelihood function.(*35, 38*) The position to cut the TCSPC histogram is determined relative to the maximum of the sum of the TCSPC histograms with a number of photons in the centre 50 % (0.3 ns after the maximum by default). There is the option to perform the lifetime fit multiple times with slightly varied initial conditions and only take the fit with the highest likelihood which reduces the lifetime uncertainty by the fitting. Besides the processing part of TrackAndTrace also the visualiser for analysed data was significantly extended. To display FLIM images the lifetime is encoded using a colour-map and the intensity by the brightness. To avoid cross-talk between lifetime and intensity, isoluminant and uniform colourmaps, created with *colorcet*,(*39*) are provided. The visualiser also can display histograms (normal, 2D or weighted) of the localisation data, filter the data, calculate and correct drift correction with Redundant Cross-Correlation (RCC),(*23*) and reconstruction of super-resolved lifetime images. The super-resolved lifetime images are calculated as weighted average of all localisations, where the weight is given by a Gaussian at the localisations position. To calculate multi-channel images, the reconstructions could be done separately by applying the corresponding filter one at a time.

### Correction of scanning artefacts

All confocal SMLM measurements in this work were done using bidirectionally scans. With the FLIMbee laser scanner, the position markers in the photon stream have an offset between the forward and the reverse scan. This causes a shift between the odd and even lines along the slow axis as shown in Fig. S6a. This shift depends on the position of the scan area, scan speed and result of an internal calibration directly before the scan. To determine the shift, an image with a pixel size along the fast axis of 5 nm is calculated and smoothed with a moving average along the fast axis. For this image, the shift between the odd and even lines that minimises the sum of the quadratic difference between each line and their adjacent lines is determined by an iterative grid search. The shift correction is done automatically by the PTU import plugin of TrackNTrace by offsetting the position markers during the generation of the photon index. Since the correction is done in the time domain, there is no loss of resolution due to interpolation.

### Image analysis with TrackNTrace framework

For the dSTORM/PAINT data analysis with TrackNTrace, the following plugins and parameters were used, as shown in the screenshot in Fig.1. The candidate detection and refinement were done with the *Cross correlation* and *TNT Fitter* plugin using the default parameters. The tracking was done with *TNT NearestNeighbor* using one as minimum track length, maximum tracking radius and no gap closing. The lifetime fitting was done with the plugin *fit lifetime*. First the position of each localisation was refitted in a sum image of each frame of the track using a Gaussian MLE fit. For the TCSPC extraction a mask radius of two and summing over all frames within the track was chosen, the fitting was done on all TCSPCs with at least 100 photons using the monoexponential MLE with a cutoff of 0.3 ns, a lifetime range of 0.3 ns to 5.5 ns and 100 fit attempts. Before the reconstruction, the localisations were filtered and only localisations with a χ^2^ of the lifetime fit between 0.9 and 1.1 were kept.

### Pattern matching analysis approach

Our pattern matching classification is based on a maximum likelihood approach previously applied to burst analysis of diffusing single molecules.(*40*) The analysis is based on the full TCSPC curve of each localisation given by {*m*(*n*)} with m denoting the number of photons in time bin *n*. The probability *P* of obtaining {*m*(*n*)} given that the localisation is of species *α* is given by

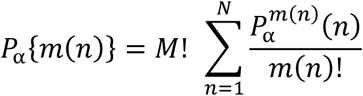

where *N* is the number of time bins, 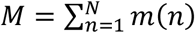, and 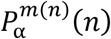 is the normalised probability distribution for species *α*. After this is done for all considered species, a molecule is classified as belonging to the species with the higher probability. To make the calculation more efficient, the logarithm of *P* is used and the term *M*! and *m*(*n*)! treated as constant, due to their independence of *α*:

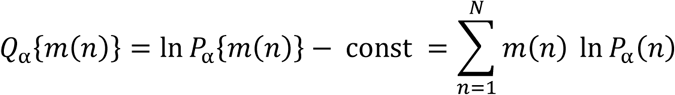

The relative probability *f*_*α*_ for species *α* is given by

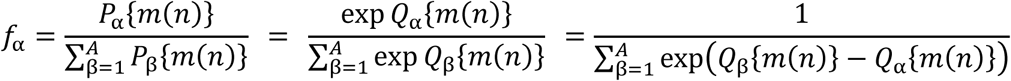

provided that a number of *A* species are considered. This relative probability *f*_*α*_ can be used to reject localisations that cannot be classified with high likelihood. A prerequisite for pattern matching is the knowledge of the TCSPC curves of each species used as references *P*_*α*_ (*n*). These were obtained by measuring a sample including only one species under the same conditions, summing the TCSPC of all localisations and normalising.

### Antibody labelling

Antibodies for staining cells, measured via lifetime-based multiplexed dSTORM imaging, were custom modified and labelled. Goat anti-rabbit IgG (Invitrogen, 31212) was used as secondary antibody for alpha-tubulin and clathrin samples. Goat anti-mouse IgG (Sigma-Aldrich, SAB3701063-1) was used as secondary antibody for beta-tubulin staining. Antibody labelling via N-hydroxysuccinimidyl esters was performed at RT for 4h in labelling buffer (100 mM sodium tetraborat (Fulka, 71999), pH 9.5) following the manufacturers standard protocol. Briefly, 100 µg antibody was reconstituted in labelling buffer using 0.5 ml spin-desalting columns (40K MWCO, ThermoFisher, 87766).

Goat-anti-rabbit IgG (Invitrogen, 31212) was modified with N-hydroxysuccinimidyl esters - PeG_4_ – trans-Cyclooctene (JenaBioscience, CLK-A137-10), using a 5 fold surplus of NHS-PeG_4_-TCO. After purification using spin-desalting columns (40K MWCO) in PBS (Sigma-Aldrich, D8537-500ML), modified goat-anti-rabbit IgG (Invitrogen, 312121) was incubated with a 10 x excess of Tetrazine-ATTO655 (ATTO-TEC, AD 655-2505) at RT and purified again after 15 min.

Goat anti-mouse IgG was incubated using a 5 x excess of N-hydroxysuccinimidyl esters – Alexa647 (LifeTech, A20106) and purified after 4 h using spin-desalting columns (40K MWCO) in PBS (Sigma-Aldrich, D8537-500ML) to remove excess dyes. Finally, antibody concentration and DOL was determined by UV-VIS absorption spectrometry (Jasco V-650).

### Cell culture and fluorescence labelling

Human mesenchymal stem cells (PT-2501) used for proof-of-principle dSTORM imaging were purchased from Lonza Group. Cells were plated on glass-bottom petri dishes (VWR) and cultured in DMEM supplemented with 10 % FCS, 2 mM L-glutamine, 1 mM sodium pyruvate, 100 U/mL penicillin-streptomycin in a humidified 5 % CO2 atmosphere at 37° C 48 hours after plating, cells were rinsed with PBS and fixed in 4 % (v/v) formaldehyde/PBS for 15 min. After that, they were permeabilized with 0.5 % Triton-X 100 in PBS for 10 min. For labelling of tubulin, permeabilized cells were blocked with 3 % BSA in PBS for 30 min at room temperature and then incubated with mouse monoclonal anti-alpha-tubulin antibody (T6199, clone DM1A, Sigma-Aldrich) diluted at 1:200 in 3 % (w/v) BSA/PBS for 1 hour. Then cells were washed in PBS, containing 0.005 % Triton X-100, incubated 1 hour with Alexa Fluor 647-conjugated cross-adsorbed goat anti-mouse IgG F(ab’)2 (A-21237, Invitrogen) diluted at 1:1000 in 3% BSA/PBS, and washed in PBS.

COS-7 cells, used for DNA-PAINT measurements, were immunostained and transfected as described previously.(*29*) Cells were stored in PBS at 4°C. Prior to immunostaining and imaging, ca. 20,000 cells/well, they were plated in 8-well chamber (155411PK, Thermo Fisher Scientific).

COS-7 cells, used for lifetime-based multiplexed dSTORM imaging, were seeded at a concentration of 2.5 × 10^4^ cells/well into 8 chambered cover glass systems with high performance cover glass (Cellvis, C8-1.5H-N) and immunostained after 3 hours of incubation at 37 °C and 5 % CO2. For microtubule and clathrin immunostaining, cells were washed with pre-warmed (37 °C) PBS (Sigma-Aldrich, D8537-500ML) and permeabilized for 2 min with 0.3 % glutaraldehyde (GA) + 0.25 % Triton X-100 (EMS, 16220 and ThermoFisher, 28314) in pre-warmed (37° C) cytoskeleton buffer (CB), consisting of 10 mM MES ((Sigma-Aldrich, M8250), pH 6.1), 150 mM NaCl (Sigma-Aldrich, 55886), 5 mM EGTA (Sigma-Aldrich, 03777), 5 mM glucose (Sigma-Aldrich, G7021) and 5 mM MgCl2 (Sigma-Aldrich, M9272). After permeabilization cells were fixed with a pre-warmed (37° C) solution of 2 % GA for 10 min. Cells were washed twice with PBS (Sigma-Aldrich, D8537-500ML). After fixation, samples were treated with 0.1 % sodium borohydride (Sigma-Aldrich, 71320) in PBS for 7 min. Cells were washed three times with PBS (Sigma-Aldrich, D8537-500ML) before blocking with 5 % BSA (Roth, #3737.3) for 30 min. Subsequently, microtubule samples were incubated with 2 ng/µl rabbit anti-alpha-tubulin primary antibody (Abcam, #ab18251) or 2 ng/µl mouse anti-beta-tubulin primary antibody (Sigma-Aldrich, T8328), the clathrin samples were incubated with 2 ng/µl rabbit anti-clathrin primary antibody (Abcam, #ab21679) in blocking buffer for 1 hour. After incubation, cells were rinsed with PBS (Sigma-Aldrich, D8537-500ML) and washed twice with 0.1 % Tween20 (ThermoFisher, 28320) in PBS (Sigma-Aldrich, D8537-500ML) for 5 min. Cells were incubated with 4 ng/µl of custom labelled goat anti-rabbit IgG – PeG_4_-ATTO655 (DOL 1.7) secondary antibodies (Invitrogen, 31212) or custom labelled goat anti-mouse IgG – Alexa647 (DOL 1.1) secondary antibodies (Sigma-Aldrich, SAB3701063-1) in blocking buffer. After secondary antibody incubation, cells were rinsed with PBS (Sigma-Aldrich, D8537-500ML) and washed twice with 0.1 % Tween20 (ThermoFisher, 28320) in PBS (Sigma-Aldrich, D8537-500ML) for 5 min. After washing, cells were fixed with 4% formaldehyde (Sigma-Aldrich, F8775) for 15 min and washed three times in PBS (Sigma-Aldrich, D8537-500ML).

### Polymer beads labelling

Polystyrene microspheres (d = 3.0-3.9 µm) coated with streptavidin (Kisker Biotech, PC-S-3.0) were labelled with fluorescent dye according to the following procedure. The solution of 500 µL water, 500 µL PBS, 100 µL of microsphere solution, and 0.5 µL of biotinylated dsDNA labelled with either Alexa 647 (1 mM in PBS) or ATTO 655 (1 mM in PBS) was centrifuged 30 min at 15 krpm. The DNA sequences are shown in the Table S1 below. The supernatant was then removed and replaced with 100 µL of PBS and the pellet was dissolved by vortexing.

50 µL of the final solution, was put on a glass coverslip and incubated for 20 min protecting it from light and evaporation. Finally, 300 µL of dSTORM imaging buffer was added on top.

## Acknowledgments

The authors thank Dr. Shama Sograte-Idrissi for COS-7 cells preparation for DNA-PAINT experiments. We are grateful to Dr. Anna Chizhik for fruitful discussions. J.E. and O.N. acknowledge financial support by the Deutsche Forschungsgemeinschaft (DFG) via project “Super-resolution microscopy through single molecule localization at cryogenic temperature” (EN297/15-1). J.C.T. is grateful to the DFG for financial support via project A05 of the SFB 937, and to the International Max Planck Research School for Physics of Biological and Complex Systems (IMPRS PBCS) for financial support. N.O. is grateful to the Deutsche Forschungsgemeinschaft (DFG) for financial support via project A10 of the SFB 803.

## Author contributions

J.C.T., D.H., R.T., M.S., O.N. and J.E. designed experiments. J.C.T. and O.N. generated and processed data. D.H. labelled antibodies and prepared cells for dSTORM measurements. E.B. prepared hMSC cell samples. N.O. and R.T. performed PAINT measurements. J.C.T. wrote the analysis software. J.C.T., D.H., N.O., R.T., M.S., O.N. and J.E wrote, edited and approved the final draft of manuscript.

## Data and materials availability

The data that support the findings of this study are available from the corresponding author upon reasonable request. The analysis software TrackNTrace is available on Github (https://github.com/scstein/TrackNTrace).

## Supporting Information

**Fig. S1.**
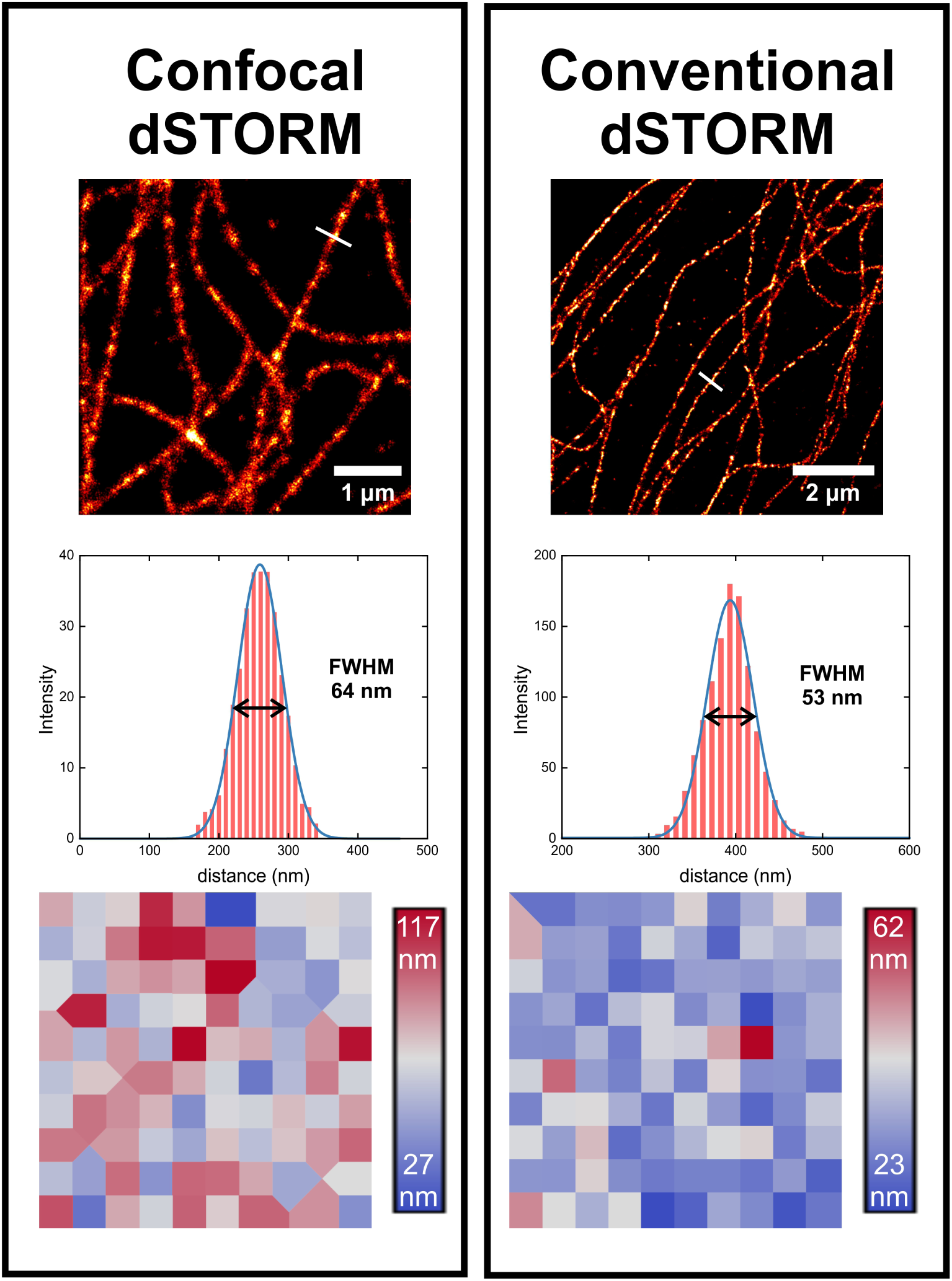
dSTORM imaging of Alexa 647 labelled tubulin in hMS cells. (**left**) Confocal dSTORM super-resolved image, intensity profile across the individual microtubule, FRC map of the corresponding region of interest. (**right**) Conventional dSTORM super-resolved image, intensity profile across the individual microtubule, FRC map of the corresponding region of interest.

**Fig. S2.**
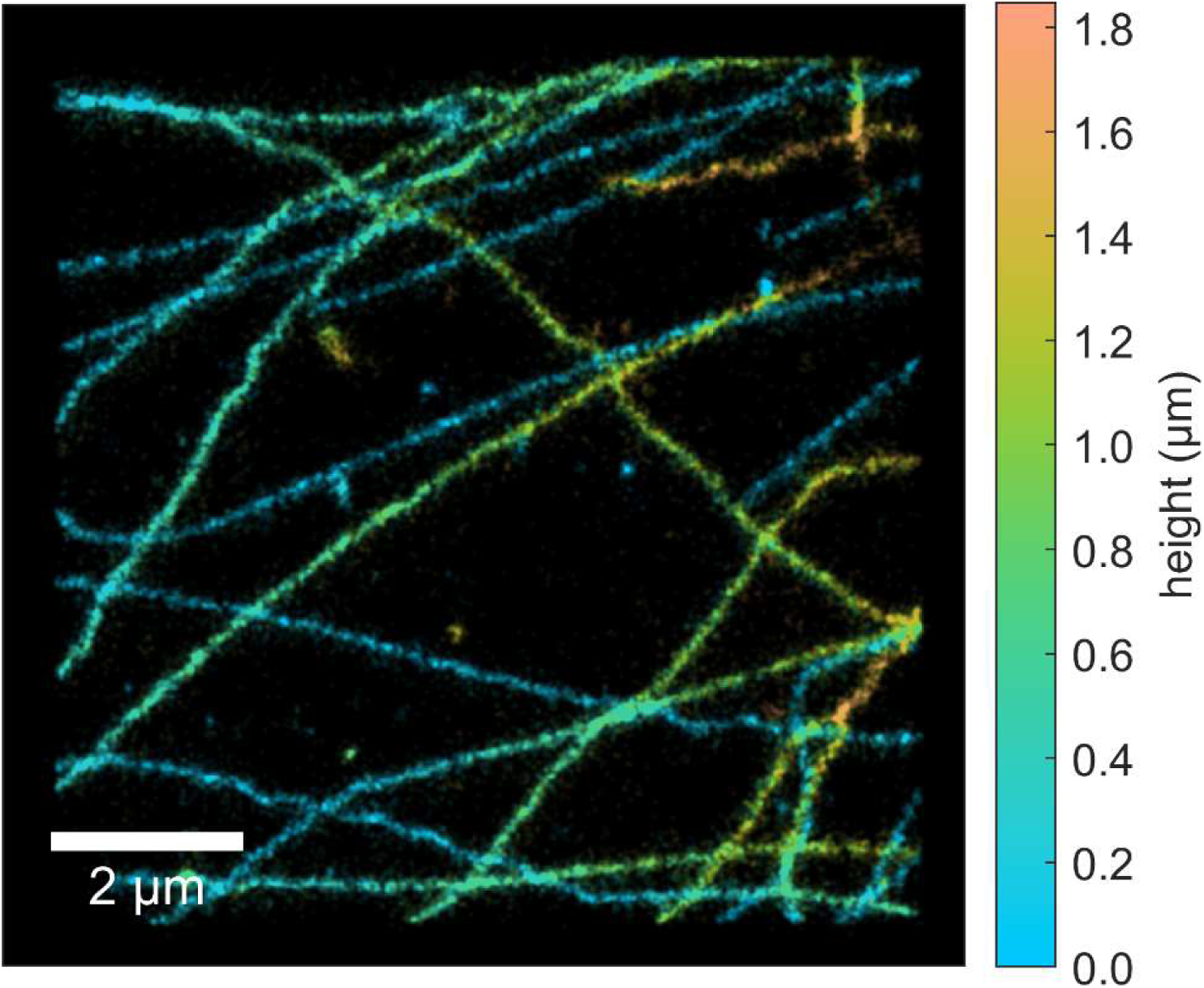
Confocal sectioning. 3D dSTORM image of Alexa647 labelled microtubules in fixed COS7 cells. The localizations are colour-coded according to their *z* position. The image was generated from the corresponding 2.1 µm *z*-stack with the step of 300 nm.

**Fig. S3.**
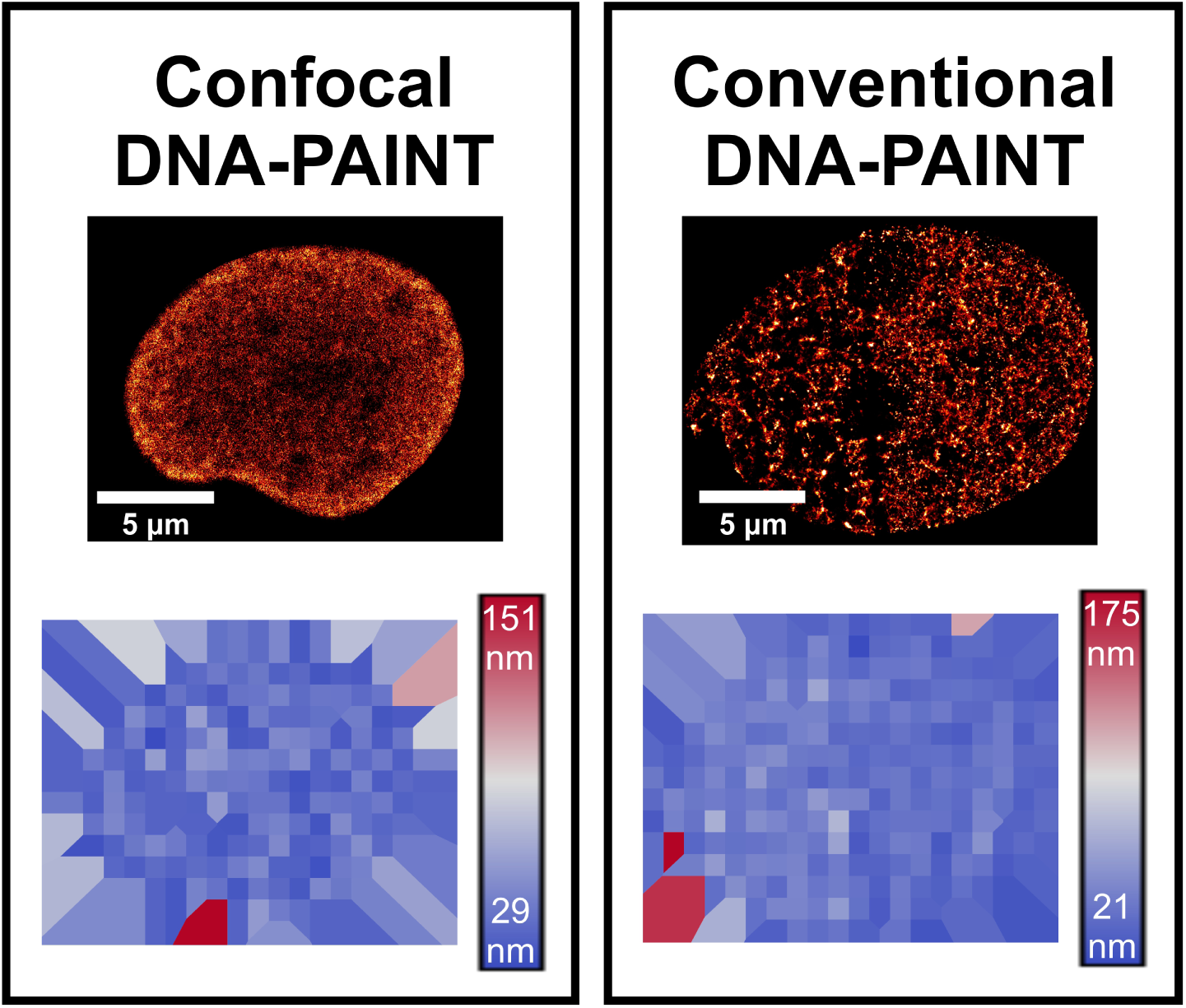
DNA-PAINT imaging of cellular chromatin in COS-7 cells utilising DNA-labelled ATTO 655. (**left**) Confocal DNA-PAINT super-resolved image and FRC map of the corresponding region of interest. (**right**) Conventional DNA-PAINT super-resolved image and FRC map of the corresponding region of interest.

**Fig. S4.**
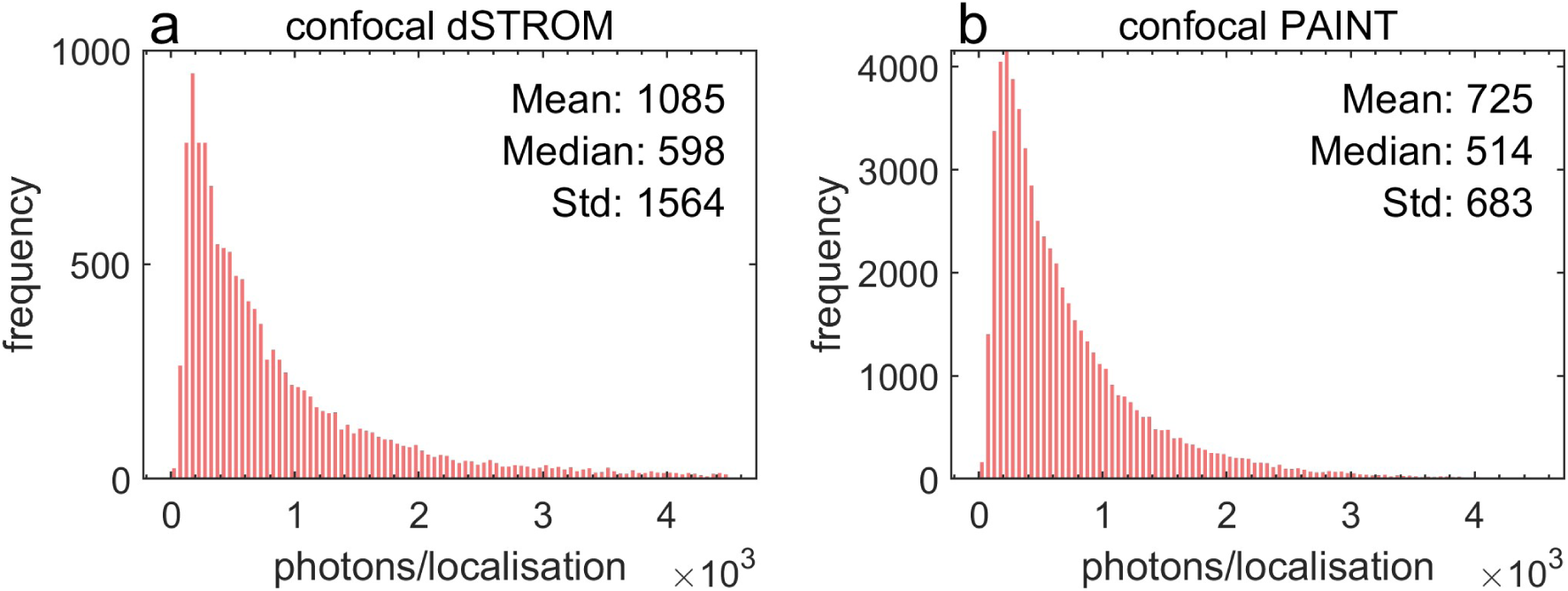
Histograms of detected photons for cell measurements shown in Fig. 2. **(a)** Confocal dSTORM utilizing Alexa 647 labelled secondary antibodies. **(b)** Confocal DNA-PAINT utilising DNA-labelled ATTO 655. The given number of photons is based on the amplitude and width of the fitted Gaussian above the background.

**Table S1.**
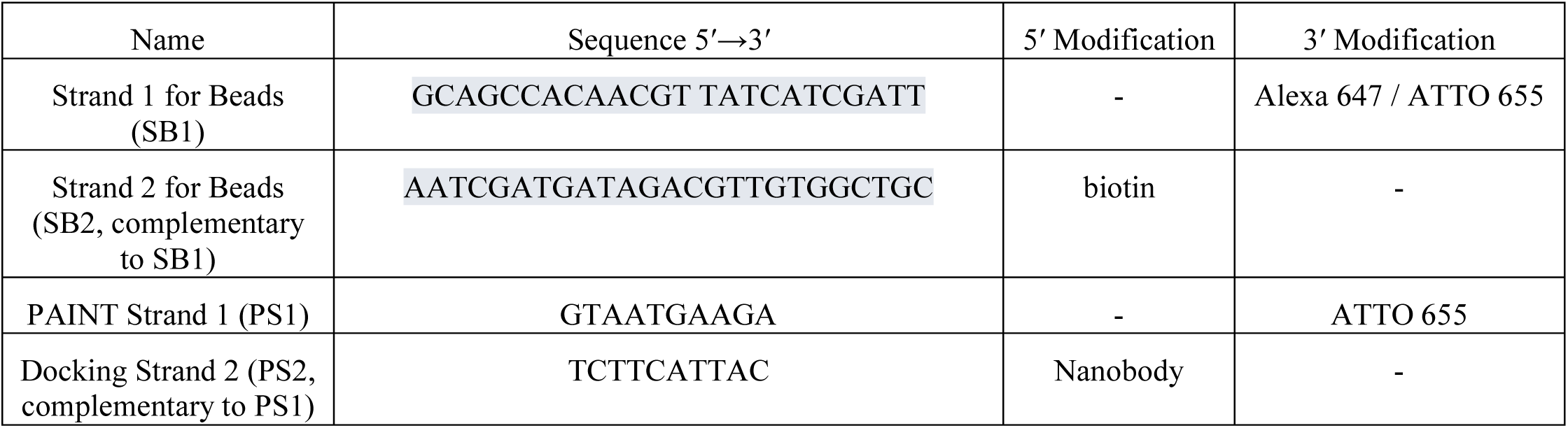
DNA sequences with modifications.

**Fig. S5.**
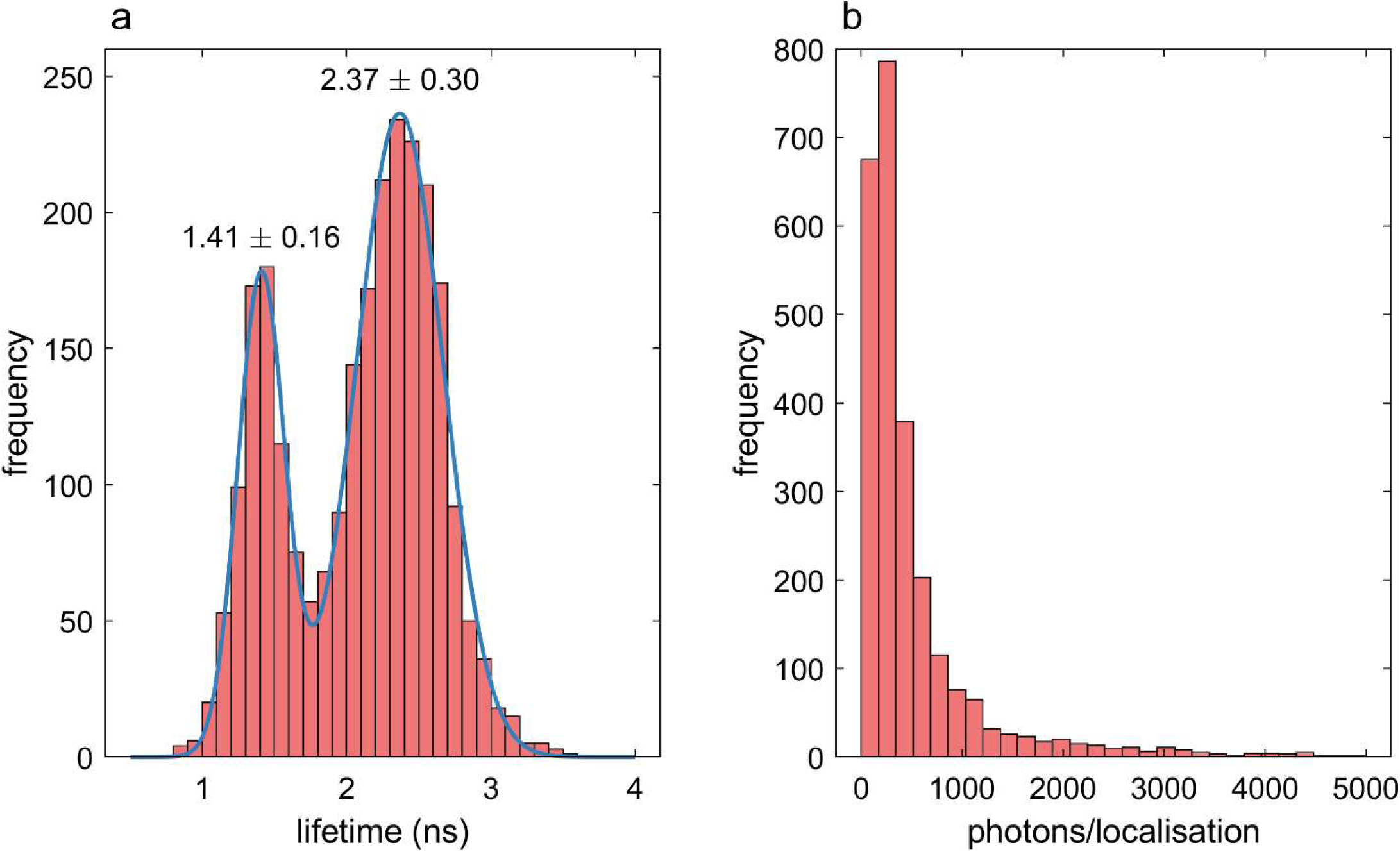
Histograms corresponding to the localisations in the bead measurement shown in Fig. 3. **(a)** Histogram of the intensity weighted fluorescence lifetimes. **(b)** Histogram of the number of photons.

**Fig. S6.**
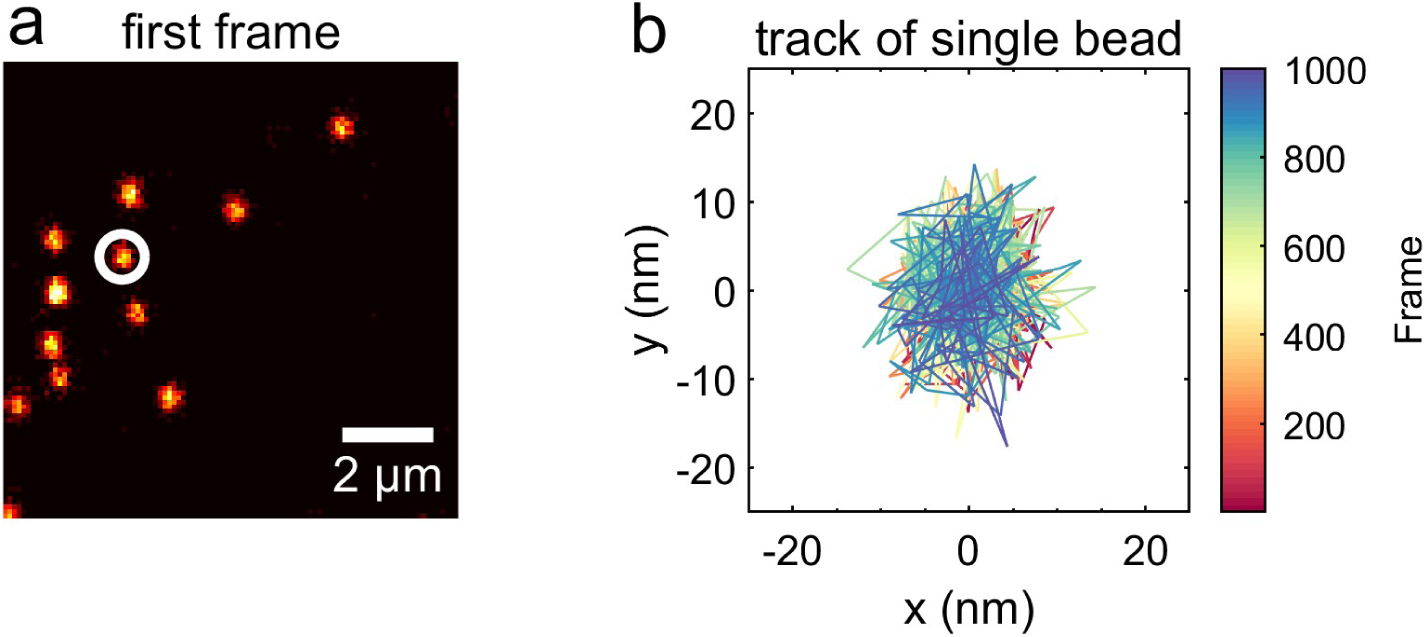
Scanning confocal measurement of TetraSpeck microspheres. **(a)** First frame of the stack. **(b)** Drift corrected position of the highlighted bead.

**Fig. S7.**
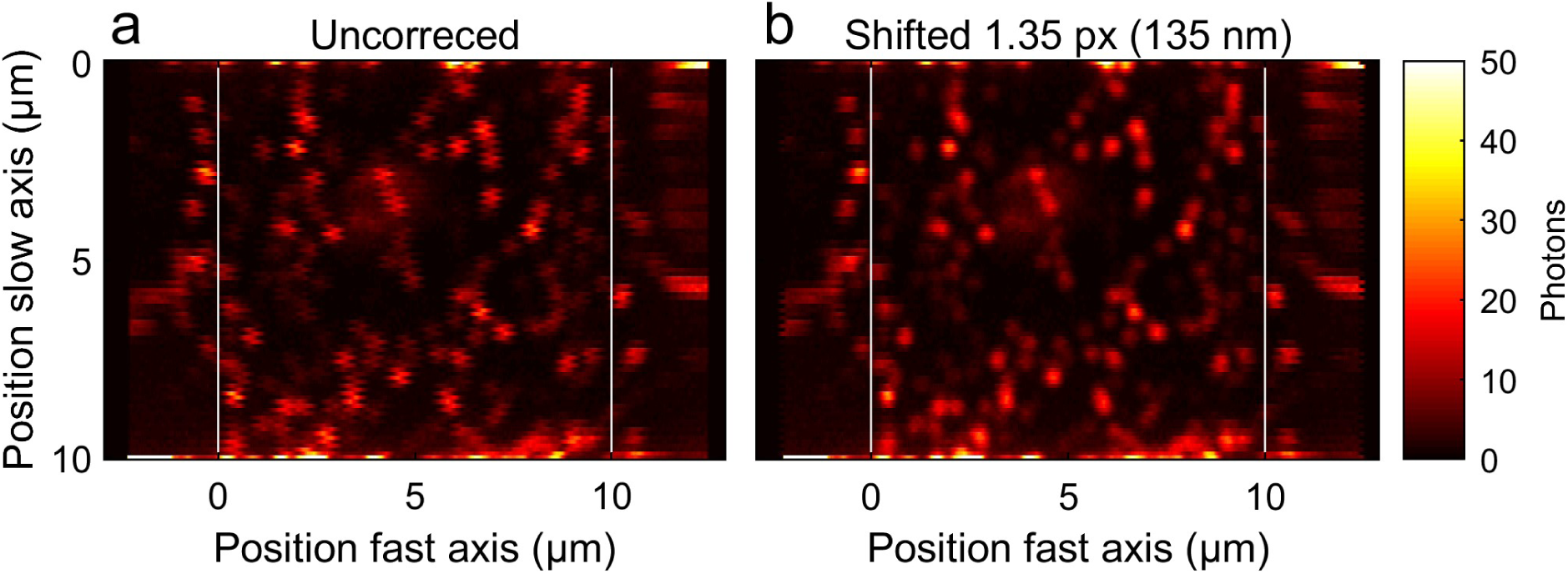
Correction of line mismatch between forward and reverse scan. **(a)** Uncorrected image. **(b)** Corrected image. For the localisation, the central part between the white lines is used and the image is generated with quadratic pixels.

